# An organelle inheritance pathway during polarized cell growth

**DOI:** 10.1101/2021.04.28.441793

**Authors:** Kathryn W Li, Ross TA Pedersen, Michelle S Lu, David G Drubin

## Abstract

Some organelles cannot be synthesized anew, so they are segregated into daughter cells during cell division. In *Saccharomyces cerevisiae*, daughter cells bud from mother cells and are populated by organelles inherited from the mothers. To determine whether this organelle inheritance occurs in a stereotyped manner, we tracked organelles using fluorescence microscopy. We describe a program for organelle inheritance in budding yeast. The cortical endoplasmic reticulum (ER) and peroxisomes are inherited concomitant with bud emergence. Next, vacuoles are inherited in small buds, followed closely by mitochondria. Finally, the nucleus and perinuclear ER are inherited when buds have nearly reached their maximal size. Because organelle inheritance timing correlates with bud morphology, which is coupled to the cell cycle, we tested whether organelle inheritance order is controlled by the cell cycle. By arresting cell cycle progression but allowing continued bud growth, we determined that organelle inheritance still occurs without cell cycle progression past S-phase, and that the general inheritance order is maintained. Thus, organelle inheritance follows a preferred order during polarized cell division, but it is not controlled exclusively by cell cycle signaling.

**Summary statement:** Organelles are interconnected by contact sites, but they must be inherited from mother cells into buds during budding yeast mitosis. We report that this process occurs in a preferred sequence.

## Introduction

Cell duplication via polarized cell growth presents a unique challenge to cellular organization. In contrast to isotropic growth – which can occur through expansion of existing cellular structure and organization – during polarized cell growth that leads to cell duplication, either a new cellular structure must be constructed from scratch, or existing components must be transported and rearranged within the cell in a regulated manner. Similarly, during development, neurons must sprout and elongate axons in order to properly wire the nervous system. This growth requires the coordination of the production and movement of various cellular building blocks, which is accomplished through signaling between the end of the growing axon and the cell body (Goldberg, 2003). Defects in organelle positioning within axons have been implicated in various neurological diseases including Charcot-Marie Tooth disorder (Suárez-Rivero et al., 2017).

Many organelles cannot be readily made de novo, and therefore must be trafficked into newly forming cellular structures, such as axons or yeast daughter cells during polarized growth (Nunnari and Walter, 1996; Warren and Wickner, 1996). This process is complicated by the fact that organelles are extensively interconnected through a network of membrane contact sites (Murley and Nunnari, 2016; Wu et al., 2018). These membrane contact sites have been implicated in crucial cellular processes ranging from lipid transfer between organelles to coordination of organelle division (AhYoung et al., 2015; Friedman et al., 2011; Lewis et al., 2016; Maeda et al., 2013). While organelle organization within the cytoplasm is critical for organelle function, how the ordered arrangement of organelles in the cytoplasm is maintained or reestablished as organelles are inherited during polarized cell growth remains a mystery.

To explore how directed movement of organelles is coordinated during polarized cell growth, we studied organelle inheritance in *S. cerevisiae*. This organism reproduces asexually by budding, wherein the daughter cell forms as a “bud” from the mother. A new cell is released by cytokinesis at the end of each cell cycle. Organelles and other cellular materials synthesized in the mother cell must be actively transported to the growing daughter cell. Numerous studies have investigated the molecular mechanisms that facilitate inheritance of individual organelles during *S. cerevisiae* bud growth. Most organelles, including endoplasmic reticulum (ER), peroxisomes, mitochondria, and vacuoles, are transported by processive myosin motors along actin cables that extend from the mother cell into the bud (Pruyne et al., 2004; Weisman, 2006). Only nuclear movement into the bud depends on microtubules (Huffaker et al., 1988), though myosin and actin cables also participate (Yin et al., 2000). Despite extensive investigations into organelle inheritance pathways in budding yeast, these pathways have mostly been studied individually. Therefore, how inheritance of different organelles is coordinated remains largely unexplored.

Recent research hints that organelle inheritance may occur in an ordered manner. One study found that membrane contact sites formed between mitochondria and the plasma membrane of emerging buds serve as anchoring sites for cytoplasmic dynein motors, which reel in astral microtubules to move the nucleus into the bud (Kraft and Lackner, 2017). Such a mechanism, wherein inherited mitochondria set up the machinery to ensure nucleus inheritance, suggests a preferred order of organelle inheritance. We wondered whether other organelles were also inherited in a preferred order.

We performed time-lapse imaging of five organelles during budding yeast mitosis to compare their inheritance. We report a preferred succession of organelles into growing buds that occurs in three stages, beginning with cortical ER and peroxisome inheritance during bud emergence, followed by the vacuole and mitochondria into small buds, and, finally, ending with nuclear and associated nuclear ER inheritance into large buds. Neither organelle inheritance itself nor the ordering of these three stages requires continuous cell cycle progression, although the nucleus is not inherited and the inheritance order of the mitochondria and vacuole is reversed when the cell cycle is arrested in S-phase, which normally begins around the time of bud emergence. Our data suggest that interdependent translocation or signaling pathways orthogonal to cell cycle signaling enforce order on organelle inheritance during *S. cerevisiae* polarized growth.

## Results and Discussion

To determine whether organelle inheritance follows a stereotyped order during budding yeast mitosis, we performed live-cell, 3D time-lapse imaging of five organelles. For each organelle, the time from bud emergence to organelle inheritance was measured. As established in the classic studies of Hartwell and colleagues (Culotti and Hartwell, 1971; Hartwell, 1971; Hartwell et al., 1970), bud morphology of logarithmically growing *S. cerevisiae* cells is highly correlated with cell cycle stage. We defined the start of each organelle inheritance time course as the time of bud emergence, allowing us to compare the timing of inheritance of organelles in different cells. We imaged cells using only bright field microscopy for varying time periods before collecting fluorescence time courses in order to capture both the moment of bud emergence and the organelle inheritance process at high temporal resolution and without significant photobleaching of genetically-encoded fluorophores. To mark the different organelles, yeast strains endogenously expressing C-terminal GFP fusions of proteins known to localize to the organelle of interest were used in most cases. Peroxisomes were visualized via Pex3-GFP (Huh et al., 2003), vacuoles were visualized via Vph1-GFP (Lu and Drubin, 2020), mitochondria were visualized via Cit1-GFP (Sawyer et al., 2019), and nuclei were visualized via Nup59-GFP (Madrid et al., 2006). The ER was visualized via expression of a single copy of GFP-HDEL integrated into the genome at the *TPI1* locus (Lu and Drubin, 2020). Cells also endogenously expressed an mCherry-tagged version of the contractile ring myosin Myo1 to clearly delineate the boundary between mother and daughter cells and to mark the onset of cytokinesis, when the ring begins to contract.

Our imaging data indicate that organelle inheritance can be described as occurring in 3 stages. The cortical ER, which lines the periphery of the cell, and the peroxisomes, are the earliest organelles inherited, with inheritance appearing to begin concomitantly with bud emergence (Fig. 1A-B). Peroxisomes are the most dynamic of the organelles that we imaged, and they became particularly difficult to track as the growing bud got bigger, allowing them more space to dynamically occupy. Nevertheless, they can clearly be seen entering the smallest buds observed (Fig. 1A and movie 1). Vacuoles and mitochondria are inherited slightly later in small buds, with inheritance commencing 10-20 minutes after bud emergence (Fig. 1C-D). Finally, nuclei are inherited once cells have reached the large-budded stage, ~40 minutes after bud emergence (Fig. 1E). Perinuclear ER, which is continuous with the nuclear envelope, behaves similarly to the nucleus itself (Fig. 1A).

**Figure 1:**
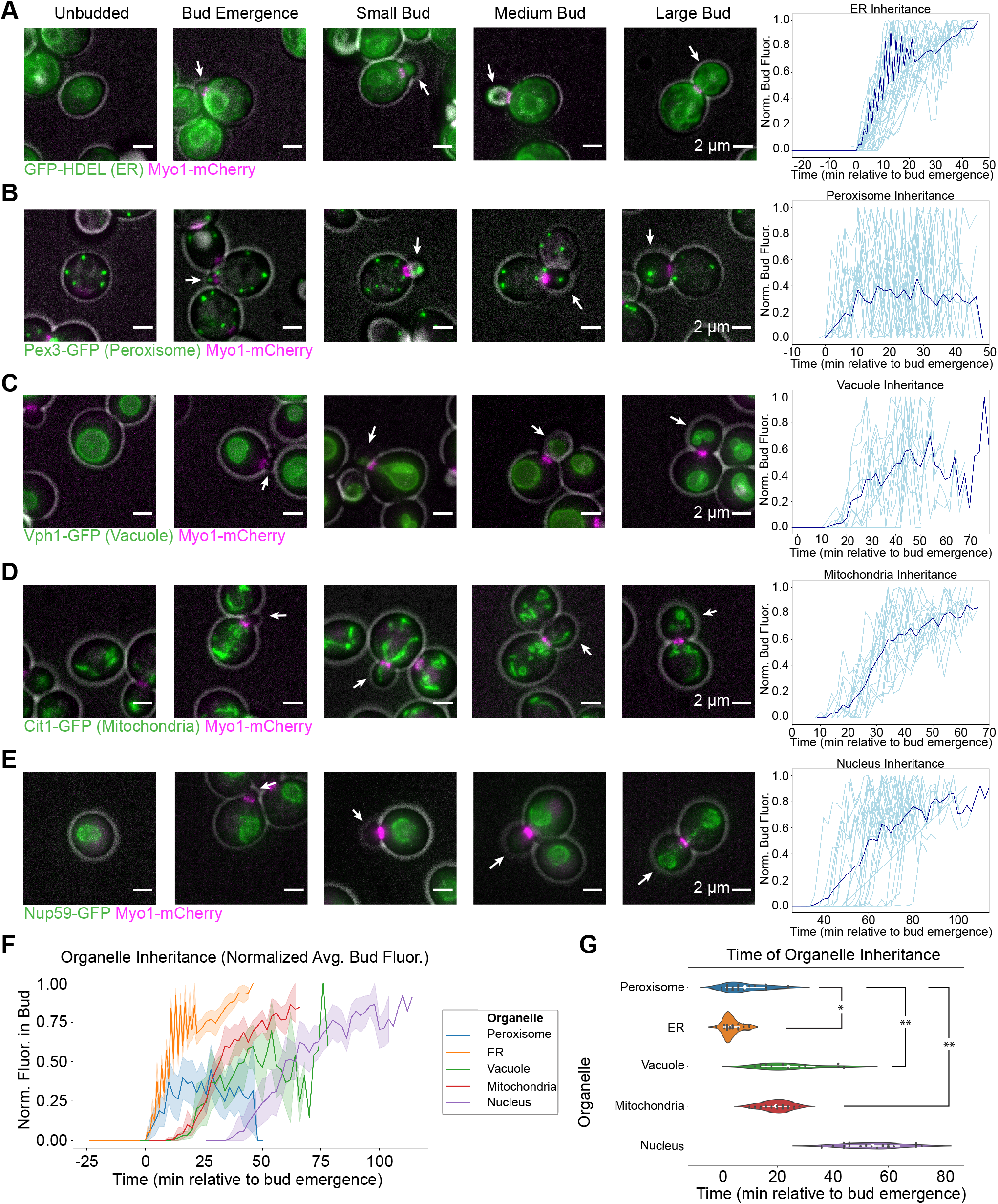
Organelle inheritance occurs in three distinct stages. (A) Left: maximum intensity projections from epifluorescence stacks of cells endogenously expressing Myo1-mCherry (magenta) to label the cytokinetic contractile ring and expressing GFP-HDEL to label the ER (green). The cell outline from bright field imaging is in gray. White arrows point to the bud in each frame. Cells at different phases of the cell cycle are juxtaposed to illustrate succession. Right: normalized fluorescence intensity of GFP-HDEL in the bud as a function of time from when bud emergence is detectable, measured in live cells. The dark blue line represents the mean fluorescence vs. time trace calculated from measurements made on 34 cells (individual measurements shown in light blue). (B-E) Left: maximum intensity projections of cells endogenously expressing Myo1-mCherry (magenta) and a Pex3-GFP peroxisome marker (green, B), Vph1-GFP vacuole marker (green, C), Cit1-GFP mitochondrial marker (green, D), or Nup59-GFP nuclear envelope marker (green, E), montaged as in (A). The cell outline from bright field imaging is in gray. Right: normalized fluorescence intensity of GFP signal in the bud as a function of time from bud emergence, measured in live cells. Dark blue lines are mean fluorescence vs. time traces calculated from 29 (B), 12 (C), 17 (D), and 22 (E) individual traces, individual measurements are shown in light blue as in (A). (F) Average fluorescence (with 95% confidence intervals) vs. time traces for organelles imaged in panels A-E plotted on the same axes for direct comparison. (G) Violin plots for the inheritance times of the organelles imaged in panels A-E with 95% confidence intervals shown in white and raw data shown as dark gray points. Inheritance time was defined as the first time after bud emergence when the bud fluorescence surpassed 0.5% of the maximum total fluorescence for the peroxisome time lapses or 2.5% of the maximum total fluorescence for the other organelle time lapses, which approximates the inflection point of the curves. Organelle inheritance times were compared by Welch’s ANOVA (F= 165) followed by Games-Howell test and asterisks indicate statistical significance between organelles whose inheritance time confidence intervals do not overlap. The 95% confidence interval for the timing of nuclear inheritance did not overlap with the 95% confidence interval for any other organelle inheritance time, so it was excluded from statistical tests and considered significantly different from all other inheritance times. * p < 0.05 (p = 0.0193), ** p < 0.01 (p = 0.0010)

Plotting the average, normalized organelle fluorescence in the bud as a function of time for all five organelles on the same axes reveals three stages of inheritance, beginning when cortical ER and peroxisomes are inherited, followed by vacuoles and mitochondria, and ending with nuclear inheritance (Fig. 1F). We functionally defined an inheritance event for an organelle as being the timepoint when fluorescence intensity for that organelle accumulated to a threshold percentage of its maximum in the bud. The threshold was defined operationally as a level of fluorescence intensity past which traces rarely fluctuated back to zero. Directly comparing the timepoint of inheritance for each organelle confirms the impression that peroxisomes and cortical ER are inherited with similar kinetics (Fig. 1G). Our statistical tests even indicated that cortical ER is inherited significantly before peroxisomes, but the difference in timing and p-value for this result were each an order of magnitude less than for all other observed differences. Mitochondria and vacuoles are inherited with similar kinetics, significantly after the peroxisomes. Finally, nuclei are inherited significantly after all other organelles. While we were able to define these three stages of inheritance by imaging organelles individually and using bud emergence as a common time reference, we were not able to resolve fine-grained differences in the timing of organelle inheritance within each stage by this analysis.

To resolve smaller differences in inheritance timing for organelles whose inheritance timing was indistinguishable using single-color imaging, we imaged pairs of organelles using two-color 3D time-lapse imaging. Although we occasionally observed that the cortical ER was inherited in emerging buds prior to peroxisomes, the inheritance timing of cortical ER and peroxisomes was still indistinguishable in the vast majority of cases (Fig. 2A, movie 1). On the other hand, when we directly compared vacuole inheritance with mitochondrial inheritance, we observed that vacuoles are inherited slightly before mitochondria (Fig. 2B, movie 2). Taken together, these results describe a timeline for organelle inheritance (Fig. 2C). Cortical ER and peroxisomes are inherited immediately upon bud emergence. Next, vacuoles and then the mitochondria are inherited at the small bud stage. Finally, nuclei are inherited at the large-budded stage.

**Figure 2:**
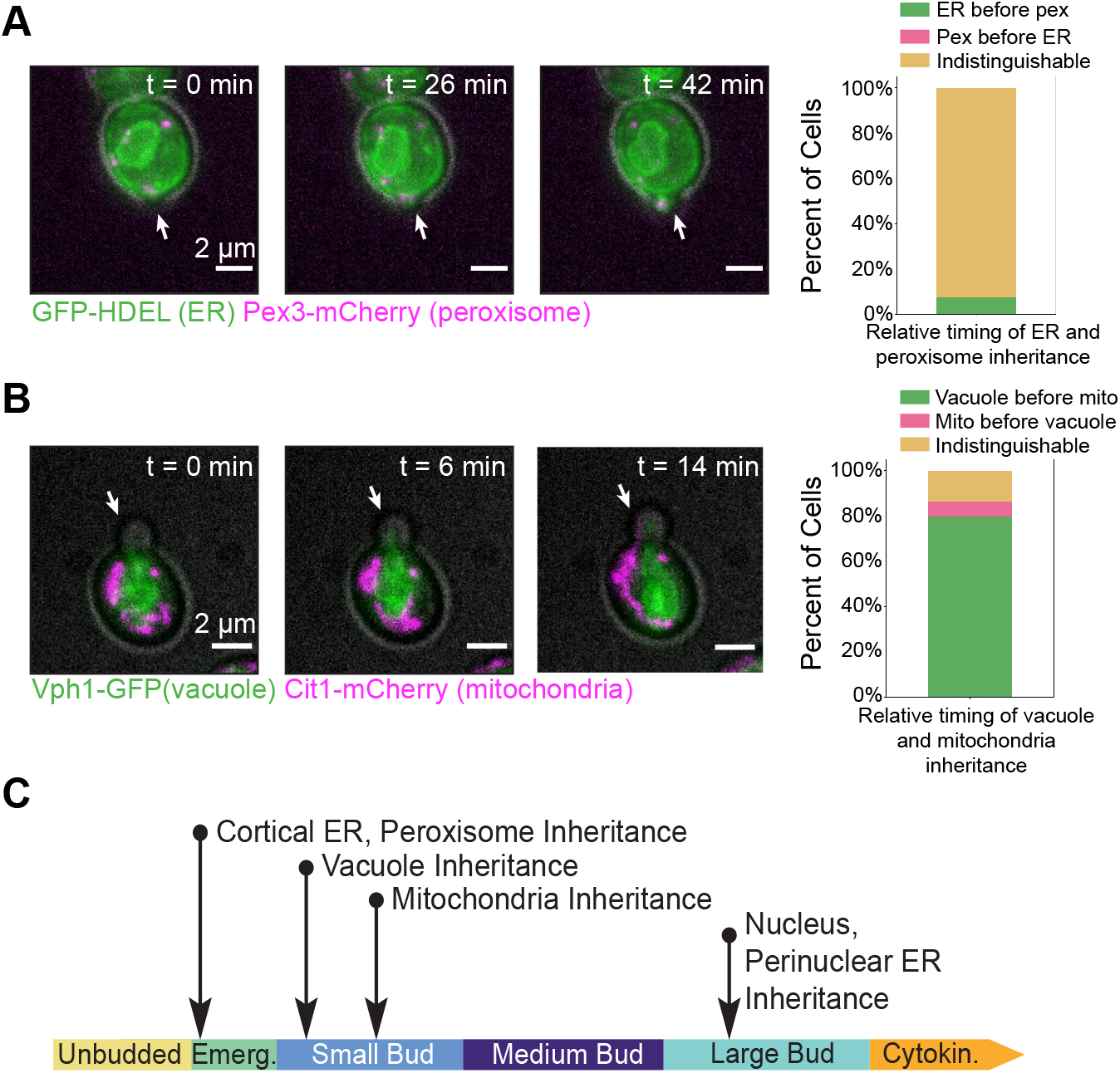
Direct comparison of inheritance within budding phases resolves order of inheritance events to elucidate an inheritance timeline. (A) Left: maximum intensity projections from a 3D time lapse epifluorescence series of a cell expressing a GFP-HDEL ER marker (green) and endogenously expressing a Pex3-mCherry peroxisome marker (magenta). The cell outline from bright field imaging is in gray. White arrows point to the bud in each frame. Minutes elapsed from the start of the time lapse are shown on the upper right corner of each frame. Right: Percent of 53 cells in which the ER is inherited before the peroxisomes (green bar), the peroxisomes are inherited before the ER (magenta bar), or the order is indistinguishable (yellow bar). (B) Left: maximum intensity projections from a 3D time lapse epifluorescence series of a cell endogenously expressing a Vph1-GFP vacuole marker (green) and a Cit1-mCherry mitochondrial marker (magenta). The cell outline from bright field imaging is in gray. Minutes elapsed from the start of the time lapse are shown on the upper right of each frame. Right: Percent of 117 cells in which the mitochondria are inherited before the vacuole (green bar), the vacuole is inherited before the mitochondria (pink bar), or the order is indistinguishable (yellow bar). (C) A timeline summarizing the observed inheritance timing of organelles during yeast budding.

We next set out to determine whether the order of organelle inheritance that we observed is coordinated with cell cycle events. Because organelle inheritance events were observed at specific points in the bud morphogenesis cycle, and because the bud morphogenesis cycle is tightly linked to the cell cycle, we wondered whether cell cycle signaling dictates the order of organelle inheritance. To test this possibility, we took advantage of the fact that hydroxyurea (HU) arrests the cell cycle of budding yeast at S-phase onset without arresting the bud morphogenesis cycle (Amberg et al., 2005). Since S-phase begins around the time of bud emergence, the result of such an arrest is progression of the bud growth cycle without corresponding cell cycle progression. The HU treatment allowed us to assess how organelles are inherited when cell cycle progression is blocked at S-phase onset.

While we hypothesized that organelle inheritance might be controlled in part by the cell cycle, we found instead that organelle inheritance mostly occurs even in the absence of cell cycle progression past S-phase onset. We arrested cells in HU for 3 hours, sufficient time for cells that were past S-phase at the time of drug addition to complete their cell cycle and arrest at the following S-phase, giving us confidence that all cells were S-phase arrested. After the 3-hour S-phase arrest, cells were morphologically arrested at the large-budded stage of the growth cycle, which normally corresponds to late M-phase (Fig. 3A). Even though cortical ER and peroxisomes are normally inherited in emerging buds (around the time of S-phase onset) and all other organelles are inherited in growing buds after S-phase onset, we nevertheless observed cortical ER, peroxisomes, vacuoles, and mitochondria in the majority of the large buds that had grown from the S-phase arrested cells (Fig. 3A-B). Nuclei, on the other hand, remained either in the mother cell (not inherited) or at the bud neck (partially inherited) (Fig. 3A-B). Thus, even in the absence of cell cycle progression past S-phase, most organelle inheritance can proceed.

**Figure 3:**
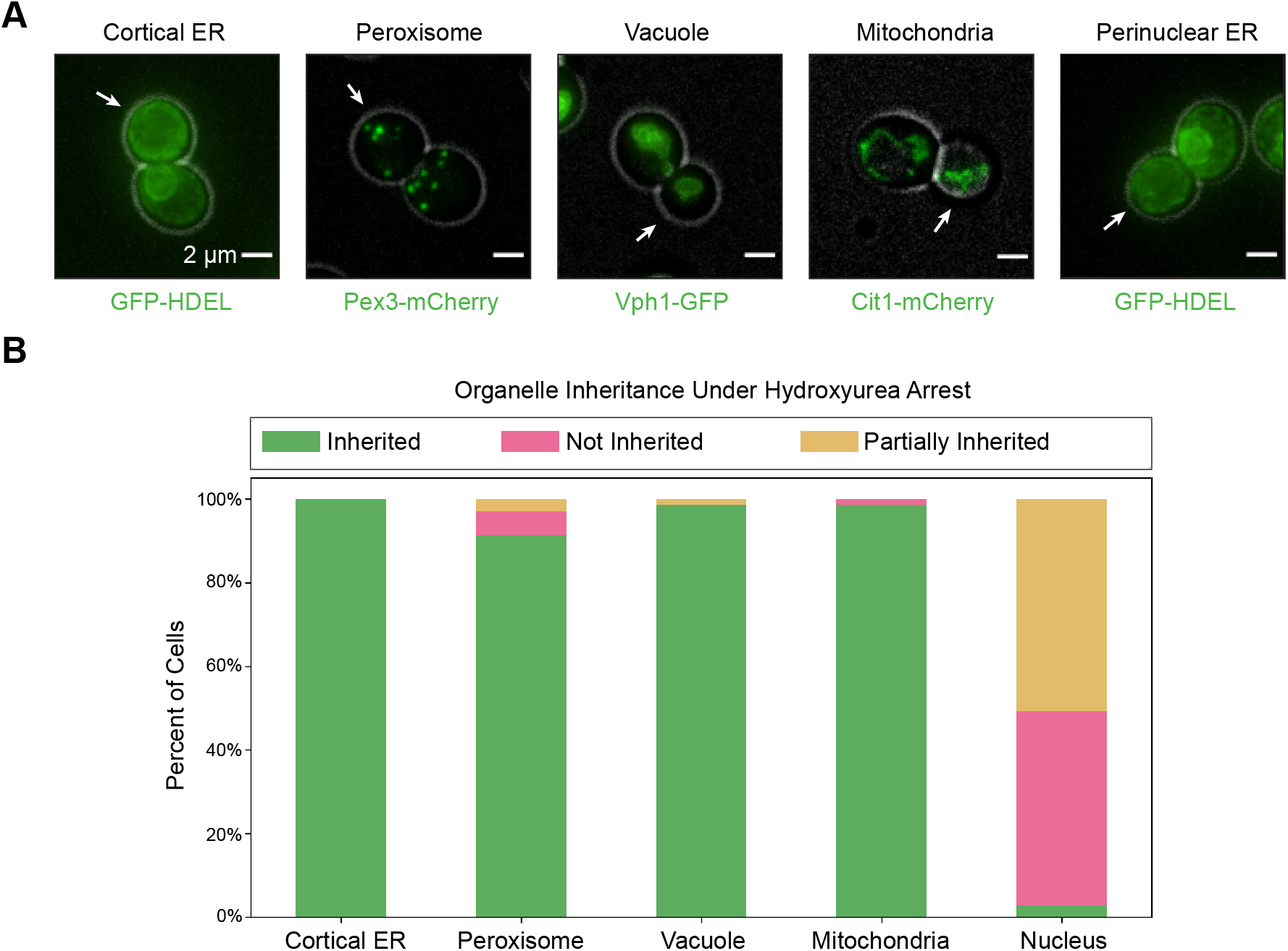
Organelle inheritance occurs in the absence of continuous cell cycle signaling. (A) Maximum intensity projections from epifluorescence stacks of cells arrested with hydroxyurea for 3 hours. From left to right, cells are expressing the ER label GFP-HDEL to visualize the cortical ER, endogenously expressing Pex3-mCherry to label peroxisomes, Vph1-GFP to label the vacuole, Cit1-mCherry to label mitochondria, and expressing the ER label GFP-HDEL to visualize the perinuclear ER (all shown in green). The cell outline from bright field imaging is in gray. White arrows point to the bud in each frame. (B) Percentage of cells (n=69 for cortical ER, peroxisome, and perinuclear ER; n=75 for vacuole and mitochondria) in which the organelle of interest was inherited (green bar), not inherited (pink bar), or partially inherited (yellow bar) after 3 hours of hydroxyurea arrest.

When we examined organelle inheritance timing, we found that the order of the three stages of inheritance we observed previously remained the same even without continuous cell cycle progression. To study organelle inheritance timing, we used alpha factor to first synchronize cells in G1, prior to S-phase and bud emergence, and then released them into HU for imaging (Amberg et al., 2005). This eliminated the possibility that bud growth observed represented cells that were past S-phase at the time of HU addition, ensuring that all bud growth occurred under HU arrest. This procedure allowed us to record a time series of organelle inheritance while bud growth was occurring in cell cycle arrested cells. We found that both the cortical ER and peroxisomes were still inherited at bud emergence, with the inheritance timing between these two organelles still mostly indistinguishable (Fig. 4A, movie 3). As in our earlier results, peroxisomes were still clearly inherited before the mitochondria, indicating that the first two stages of organelle inheritance that we had observed were still separable (Fig. 4B, movie 4). In a departure from our results with unmanipulated cells, we observed the mitochondria being inherited before the vacuole, but both organelles were still inherited into small buds (Fig. 4C, movie 5). Thus, despite small changes in the order of organelle inheritance within a given stage, such as with the vacuole and mitochondria, the overall order of the different stages remained the same in the absence of cell cycle progression past S-phase.

**Figure 4:**
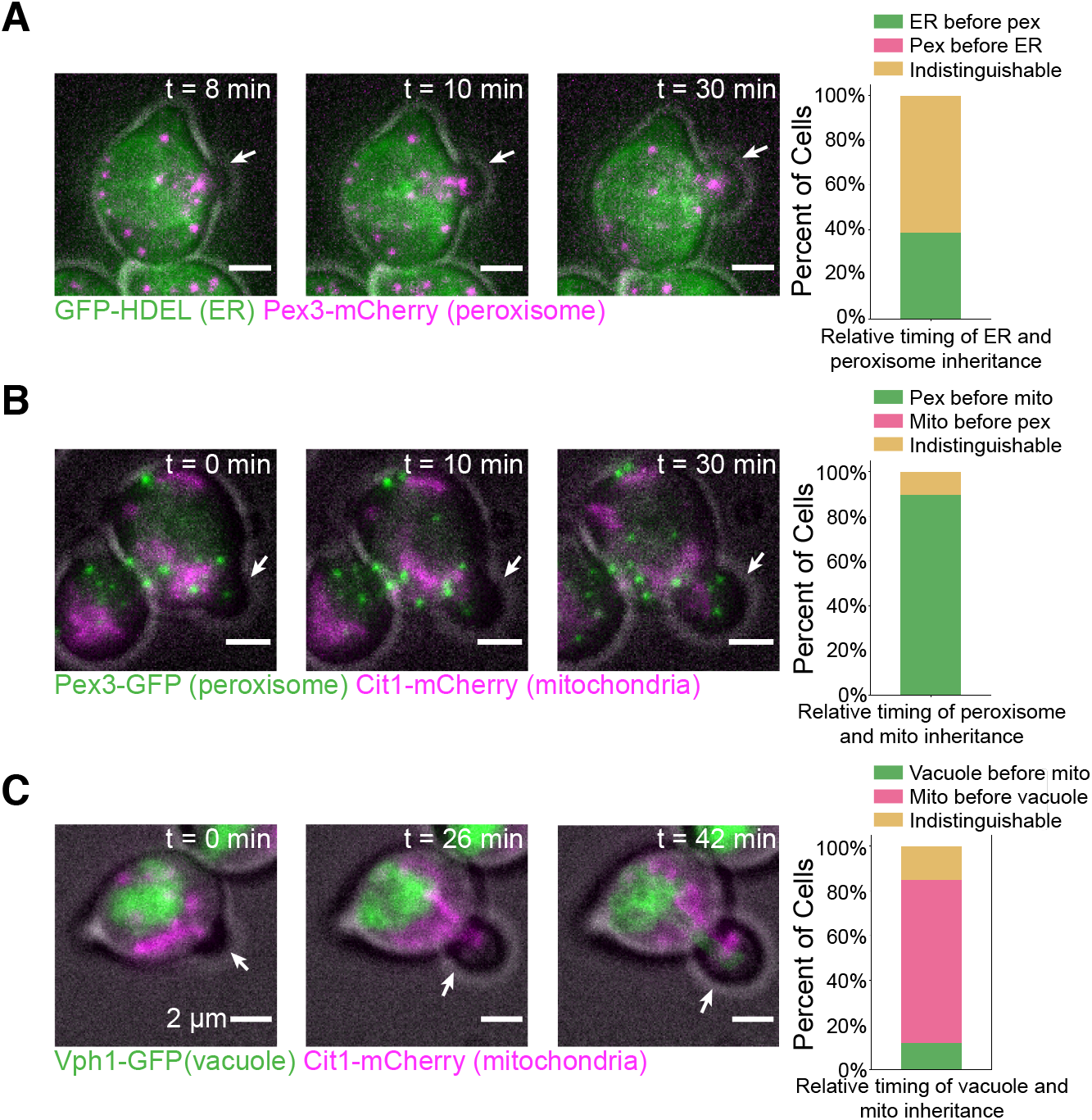
Order of organelle inheritance remains largely normal without cell cycle progression past S-phase. All images on the left show maximum intensity projections from 3D epifluorescence time lapse series of cells after arresting with alpha factor for 3 hours and releasing into hydroxyurea. The cell outline from bright field imaging is in gray. White arrows point to the bud in each frame. Minutes elapsed from the start of the time lapse are shown on the upper right of each frame. (A) Left: A cell expressing GFP-HDEL (green) and endogenously expressing Pex3-mCherry (magenta). Right: Percentage of 31 cells where the ER is inherited before the peroxisomes (green bar), the peroxisomes are inherited before the ER (magenta bar), or the exact order is indistinguishable (yellow bar). (B) Left: A cell endogenously expressing Pex3-GFP (green) and Cit1-mCherry (magenta). Right: Percent of 38 cells in which the peroxisomes are inherited before the mitochondria (green bar), the mitochondria is inherited before the peroxisomes (magenta bar), or the order is indistinguishable (yellow bar). (C) Left: A cell endogenously expressing Vph1-GFP (green) and Cit1-mCherry (magenta). Right: Percent of 38 cells in which the vacuole is inherited before the mitochondria (green bar), the mitochondria are inherited before the vacuole (magenta bar), or the order is indistinguishable (yellow bar).

Our results demonstrate that organelle inheritance in budding yeast occurs in a predictable order. Previous studies of the molecular mechanisms underlying organelle inheritance in this organism typically studied organelles individually, going so far as to demonstrate that failed inheritance of one organelle had no major effects on the inheritance of others (see for example: Du *et al.*, 2001; Ishikawa *et al.*, 2003). More recent studies, however, hint that some organelle inheritance pathways are interdependent on one another (Kraft and Lackner, 2017). The fact that organelle inheritance follows a stereotyped timeline (Fig. 2C) suggests that other such interdependent organelle inheritance pathways may be at play during budding yeast mitosis.

We also found that most organelles are inherited in the absence of cell cycle events subsequent to entry into S-phase. Some studies have shown that proteins involved in inheritance of specific organelles may be regulated by cell cycle signaling (Fagarasanu et al., 2005; Peng and Weisman, 2008). However, our results demonstrate that successful inheritance of the cortical ER, peroxisomes, vacuoles, and mitochondria still occurs under S-phase arrest (Fig. 3A-B). Moreover, the coupling of organelle inheritance to bud morphology remains largely unchanged, with organelles being inherited during the same morphological stages as described in the timeline (Fig. 2C). This observation suggests that while cell cycle signaling may influence inheritance of individual organelles, different signaling pathways regulate the relative order in which organelles are inherited. Given that inheritance of each organelle occurs at distinct stages of bud emergence or growth, the timing of organelle inheritance may be in part coordinated with bud morphogenesis. A recent study described how non cell cycle cues – including signaling by the polarity regulator Cdc42, priming of septins, and cell wall weakening – control the timing of bud emergence (Lai et al., 2018). Furthermore, one study showed that loss of cortical ER inheritance disrupts septin assembly, hinting that organelle inheritance and bud morphogenesis may be interdependent (Loewen et al., 2007). After bud emergence, inheritance of organelles may be governed by interdependent inheritance pathways. These pathways may ensure that organelle-organelle contact sites and their associated inter-organelle functions, such as lipid exchange, are maintained after cytokinesis.

## Materials and Methods

### Strains and Plasmids

All strains used in this study are listed in Table S1. Budding yeast strains were all derived from wild-type diploid DDY1102 and propagated using standard techniques (Amberg *et al.*, 2005). The GFP-HDEL strain was constructed by integrating a GFP-HDEL::LEU plasmid (courtesy of Laura Lackner) at the *TPI1* locus. This plasmid contains the pRS305 backbone and contains the *TPI1* promoter followed by the leader sequence of *KAR2* (a.a. 1-52), followed by GFP, and then HDEL. C-terminal GFP and mCherry fusions were constructed as described previously (Lee et al., 2013; Longtine et al., 1998) and verified using PCR.

### Live-Cell Imaging

Cells grown to mid-log phase in imaging media (synthetic minimal media supplemented with adenine, L-histidine, L-leucine, L-lysine, L-methionine, uracil, and 2% glucose) were immobilized on coverslips coated with 0.2 mg/ml concanavalin A.

Epifluorescence microscopy was conducted using a Nikon Eclipse Ti inverted microscope with a Nikon 100× 1.4-NA Plan Apo VC oil-immersion objective and an Andor Neo 5.5 sCMOS camera. A Lumencore Spectra X LED light source with an FF-493/574-Di01 dual-pass dichroic mirror and FF01-512/630-25 dual-pass emission filters (Semrock) was used for two-color imaging of GFP and mCherry channels. This setup was controlled by Nikon Elements software. Imaging was conducted in a room maintained at 23-25°C.

To study organelle inheritance events relative to time of bud emergence, cells were first imaged under bright field for various times to capture the moment bud emergence occurred. Immediately afterwards, cells were imaged using epifluorescence microscopy to monitor inheritance of fluorescently labelled organelles.

Image visualization was carried out with Fiji software (National Institutes of Health). For figure panels, cells were cropped, background signal was uniformly subtracted, and photobleaching was corrected using a custom Fiji macro. Figures were then assembled in Adobe Illustrator 2019.

### Hydroxyurea and alpha factor experiments

Appropriate working concentrations of hydroxyurea and alpha factor were determined empirically (Fig. S1A-B) and were generally in line with concentrations used previously (Amberg et al., 2005).

Hydroxyurea was purchased from Sigma-Aldrich. For single arrest experiments, cells were adhered to coverslips with concanavalin A and treated with 500 μL of 200 μM hydroxyurea in imaging media for three hours. Cells were then imaged using epifluorescence microscopy.

Alpha factor was synthesized by David King (University of California, Berkeley) and stored as a stock at 10 mg/mL in 0.1 M Sodium Acetate buffer (pH 5.2). Cells were adhered to coverslips with concanavalin A and submerged in 500 μL of 3 μM alpha factor in imaging media for three hours. To release from the arrest, the imaging media with alpha factor was removed and new media with 0.1 mg/ml Pronase E (Sigma P-6911) was added to inactivate any remaining alpha factor. This process was repeated 2-3 times after which 1.5 mL of imaging media with 300 μM hydroxyurea was added.

### Data analysis

To measure fluorescence intensity of an organelle in the bud during inheritance, time lapse images of cells at the appropriate bud growth stage were cropped. Cropped time lapses were segmented using the Allen Cell Structure Segmenter (Allen Institute for Cell Science, Seattle, WA) and stacks of segmented images at each time point were converted to summed projections. These time lapses were then analyzed using Fiji software (National Institutes of Health). Raw integrated fluorescence intensity was measured in manually drawn selections surrounding and encompassing the bud, and normalized relative to the maximum total fluorescence for each time lapse. Time relative to bud emergence was calculated using the corresponding bright field time lapse.

Cells were first visualized in Fiji and background subtraction and photobleaching correction were applied as described in Live-Cell imaging. For the hydroxyurea-only arrest experiments, organelles in cells were characterized as “inherited” if they were clearly present in the bud at the time of imaging, “not inherited” if no organelles were seen in the bud, and “partially inherited” if all organelles were either in the mother cell or crossing the bud neck. In characterizing the relative order of inheritance for two organelles, one organelle was considered inherited first if during the time lapse the organelle entered the bud before the other organelle or if the organelle was present in the bud before the other organelle began to be segregated to the bud. The order was considered “indistinguishable” if both organelles appeared to be inherited at the same time.

### Statistics and reproducibility of experiments

All data presented were replicated in at least three distinct experiments. Multiple cells from each replicate were analyzed and data from different days were pooled together because they were indistinguishable. The number of cells analyzed at each timepoint for Figure 1 is displayed in Figure S2, and the number of cells analyzed for the remainder of the results is shown in the figure legend.

Statistical analyses (Welch’s ANOVA test followed by Games-Howell posthoc test) were performed in Python using the Pingouin statistical package (Vallat, 2018).

## Acknowledgements

We thank Cyna Shirazinejad for data analysis assistance and Zane Bergman and Jonathan Wong for advice on experimental design. This work was funded by NIGMS grant <u>R35 GM118149</u> to DGD.

## Figures and tables

**Figure S1:**
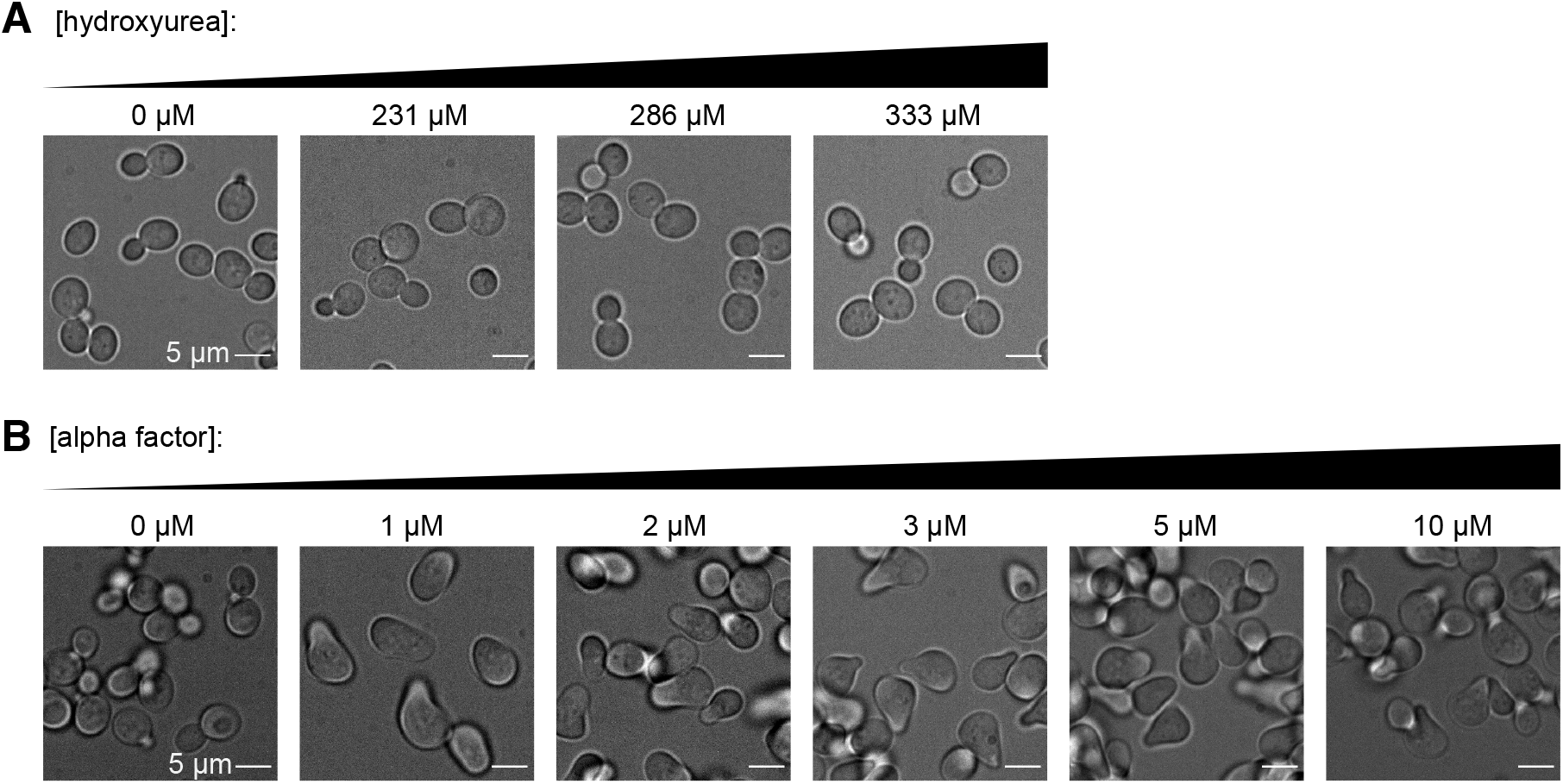
Titrations of hydroxyurea and mating factor on organelle-labelled cells. (A) Effects of hydroxyurea on cells of our background. Cells were treated with hydroxyurea for three hours at the indicated concentration and imaged in brightfield. (B) Effects of alpha factor on matA cells of our background. Cells were submerged in alpha factor for four hours at the indicated concentration and imaged in brightfield.

**Figure S2:**
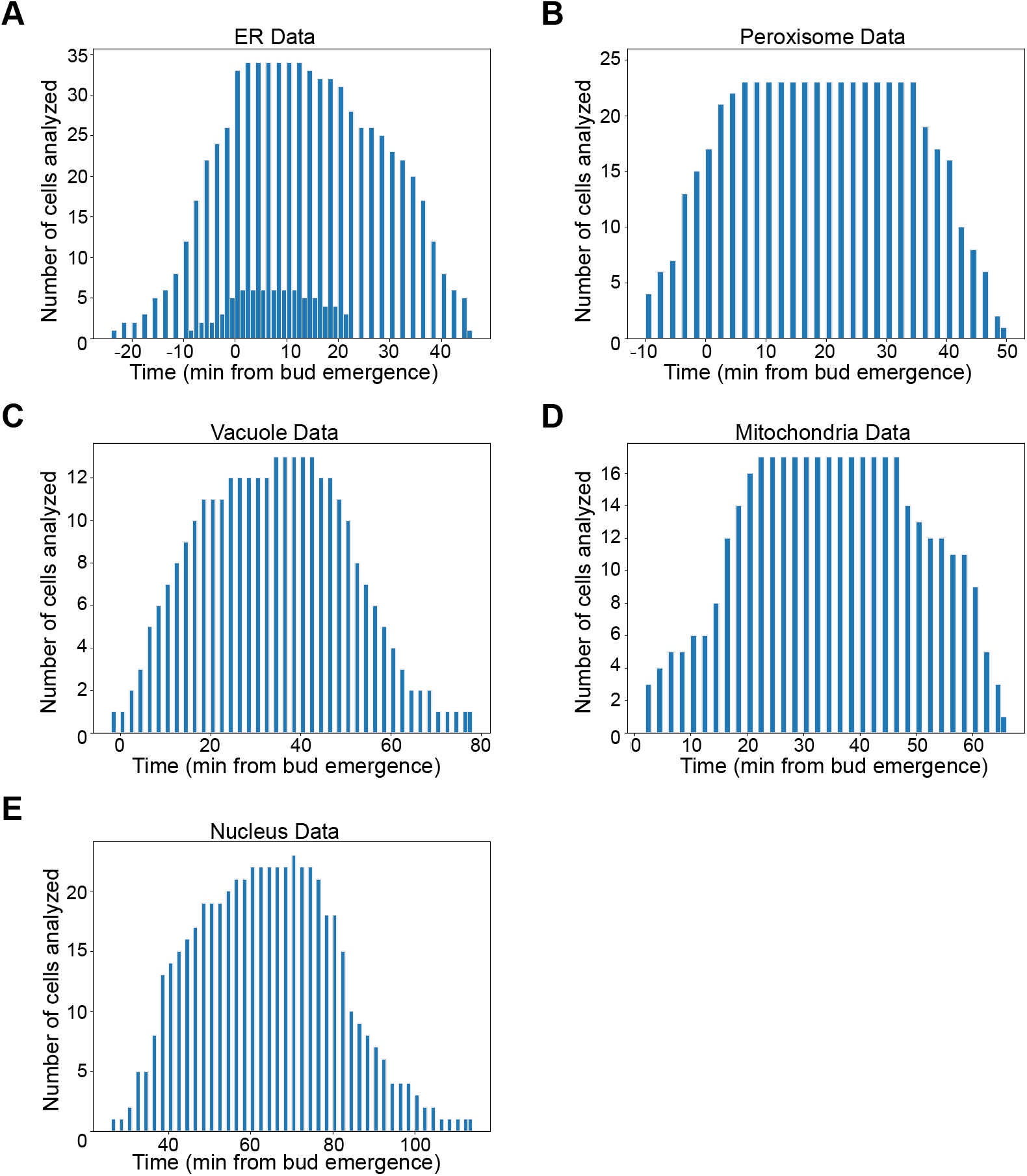
Distributions of data points used in graphing organelle inheritance. Histograms depicting the number of cells analyzed per time point past bud emergence used to plot normalized bud fluorescence in Figure 1F. Data from cells labelling the ER (A), peroxisomes (B), vacuoles (C), mitochondria (D), and nucleus (E) are shown in separate panels.

**Movie 1: Inheritance of ER and peroxisomes into emerging buds**

Maximum intensity projection movie from 3D time lapse epifluorescence imaging of a cell expressing a GFP-HDEL ER marker (green) and endogenously expressing a Pex3-mCherry peroxisome marker (magenta). The cell outline from bright field imaging is in gray.

**Movie 2: Inheritance of vacuoles and mitochondria into small buds**

Maximum intensity projection movie from 3D time lapse epifluorescence imaging of a cell endogenously expressing a Vph1-GFP vacuole marker (green) and a Cit1-mCherry mitochondrial marker (magenta). The cell outline from bright field imaging is in gray.

**Movie 3: Inheritance of ER and peroxisomes in hydroxyurea-arrested cells**

Maximum intensity projection movie from 3D time lapse epifluorescence imaging of a cell expressing a GFP-HDEL ER marker (green) and endogenously expressing a Pex3-mCherry peroxisome marker (magenta), arrested in early S-phase with hydroxyurea. The cell outline from bright field imaging is in gray.

**Movie 4: Inheritance of peroxisomes and mitochondria in hydroxyurea-arrested cells**

Maximum intensity projection movie from 3D time lapse epifluorescence imaging of a cell endogenously expressing a Pex3-GFP peroxisome marker (green) and a Cit1-mCherry mitochondrial marker (magenta), arrested in early S-phase with hydroxyurea. The cell outline from bright field imaging is in gray.

**Movie 5: Inheritance of peroxisomes and mitochondria in hydroxyurea-arrested cells**

Maximum intensity projection movie from 3D time lapse epifluorescence imaging of a cell endogenously expressing a Pex3-GFP peroxisome marker (green) and a Cit1-mCherry mitochondrial marker (magenta), arrested in early S-phase with hydroxyurea. The cell outline from bright field imaging is in gray.

**Table 1.**
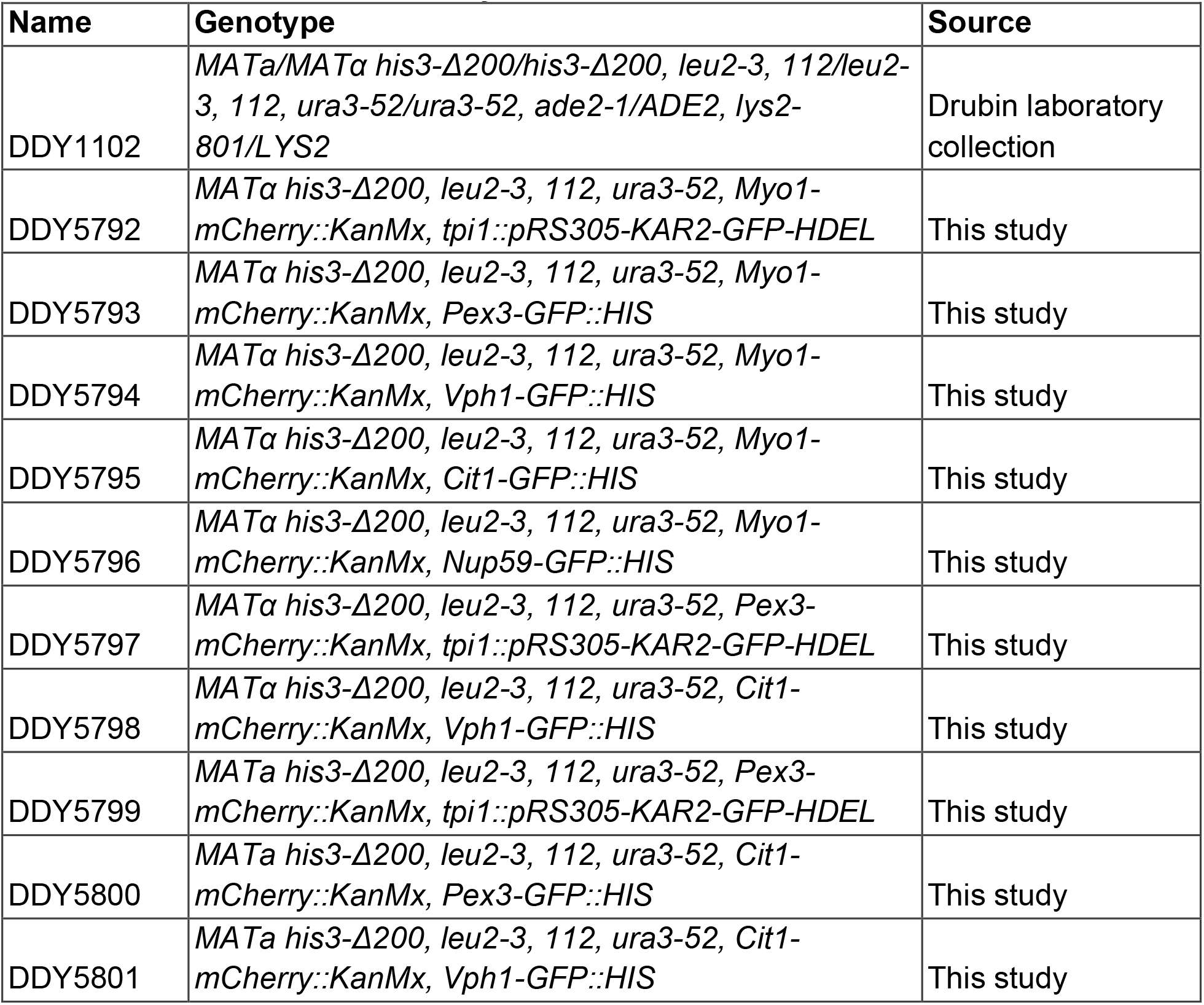
Strains used in this study

## References

AhYoung, A. P., Jiang, J., Zhang, J., Dang, X. K., Loo, J. A., Zhou, Z. H. and Egea, P. F. (2015). Conserved SMP domains of the ERMES complex bind phospholipids and mediate tether assembly. Proc. Natl. Acad. Sci. U. S. A. 112, E3179–E3188.

Amberg, D. C., Burke, D. J. and Strathern, J. N. (2005). Methods in Yeast Genetics: A Cold Spring Harbor Laboratory Course Manual, 2005 Edition.

Culotti, J. and Hartwell, L. H. (1971). Genetic control of the cell division cycle in yeast. 3. Seven genes controlling nuclear division. Exp. Cell Res. 67, 389–401.

Du, Y., Pypaert, M., Novick, P. and Ferro-Novick, S. (2001). Aux1p/Swa2p is required for cortical endoplasmic reticulum inheritance in Saccharomyces cerevisiae. Mol. Biol. Cell 12, 2614–2628.

Fagarasanu, M., Fagarasanu, A., Tam, Y. Y. C., Aitchison, J. D. and Rachubinski, R. A. (2005). Inp1p is a peroxisomal membrane protein required for peroxisome inheritance in Saccharomyces cerevisiae. J. Cell Biol. 169, 765–775.

Friedman, J. R., Lackner, L. L., West, M., DiBenedetto, J. R., Nunnari, J. and Voeltz, G. K. (2011). ER tubules mark sites of mitochondrial division. Science 334, 358–362.

Goldberg, J. L. (2003). How does an axon grow? Genes Dev. 17, 941–958.

Hartwell, L. H. (1971). Genetic Control of the Cell Division Cycle in Yeast II. Genes Controlling DNA Replication and its Intiation. J. Mol. Biol. 59, 183–194.

Hartwell, L. H., Culotti, J. and Reid, B. (1970). Genetic control of the cell-division cycle in yeast. I. Detection of mutants. Proc. Natl. Acad. Sci. U. S. A. 66, 352–359.

Huffaker, T. C., Thomas, J. H. and Botstein, D. (1988). Diverse effects of β-tubulin mutations on microtubule formation and function. J. Cell Biol. 106, 1997–2010.

Huh, K., W., Falvo, V., J., Gerke, C., L., Carroll, S., A., Howson, W., R., et al. (2003). Global analysis of protein localization in budding yeast. Nature 425, 686–691.

Ishikawa, K., Catlett, N. L., Novak, J. L., Tang, F., Nau, J. J. and Weisman, L. S. (2003). Identification of an organelle-specific myosin V receptor. J. Cell Biol. 160, 887–897.

Kraft, L. M. and Lackner, L. L. (2017). Mitochondria-driven assembly of a cortical anchor for mitochondria and dynein. J. Cell Biol. 216, 3061–3071.

Lai, H., Chiou, J. G., Zhurikhina, A., Zyla, T. R., Tsygankov, D. and Lew, D. J. (2018). Temporal regulation of morphogenetic events in Saccharomyces cerevisiae. Mol. Biol. Cell 29, 2069–2083.

Lee, S., Lim, W. A. and Thorn, K. S. (2013). Improved Blue, Green, and Red Fluorescent Protein Tagging Vectors for *S. cerevisiae*. PLoS One 8, 4–11.

Lewis, S. C., Uchiyama, L. F. and Nunnari, J. (2016). ER-mitochondria contacts couple mtDNA synthesis with Mitochondrial division in human cells. Science 353,.

Loewen, C. J. R., Young, B. P., Tavassoli, S. and Levine, T. P. (2007). Inheritance of cortical ER in yeast is required for normal septin organization. J. Cell Biol. 179, 467–483.

Longtine, M. S., McKenzie, a, Demarini, D. J., Shah, N. G., Wach, a, Brachat, a, Philippsen, P. and Pringle, J. R. (1998). Additional modules for versatile and economical PCR-based gene deletion and modification in *Saccharomyces cerevisiae*. Yeast 14, 953–61.

Lu, M. S. and Drubin, D. G. (2020). Cdc42 GTPase regulates ESCRTs in nuclear envelope sealing and ER remodeling. J. Cell Biol. 219,.

Madrid, A. S., Mancuso, J., Cande, W. Z. and Weis, K. (2006). The role of the integral membrane nucleoporins Ndc1p and Pom152p in nuclear pore complex assembly and function. J. Cell Biol. 173, 361–371.

Maeda, K., Anand, K., Chiapparino, A., Kumar, A., Poletto, M., Kaksonen, M. and Gavin, A. C. (2013). Interactome map uncovers phosphatidylserine transport by oxysterol-binding proteins. Nature 501, 257–261.

Murley, A. and Nunnari, J. (2016). The Emerging Network of Mitochondria-Organelle Contacts. Mol. Cell 61, 648–653.

Nunnari, J. and Walter, P. (1996). Regulation of organelle biogenesis. Cell 84, 389–394.

Peng, Y. and Weisman, L. S. (2008). The Cyclin-Dependent Kinase Cdk1 Directly Regulates Vacuole Inheritance. Dev. Cell 15, 478–485.

Pruyne, D., Legesse-Miller, A., Gao, L., Dong, Y. and Bretscher, A. (2004). Mechanisms of Polarized Growth and Organelle Segregation in Yeast. Annu. Rev. Cell Dev. Biol. 20, 559–591.

Sawyer, E. M., Joshi, P. R., Jorgensen, V., Yunus, J., Berchowitz, L. E. and Ünal, E. (2019). Developmental regulation of an organelle tether coordinates mitochondrial remodeling in meiosis. J. Cell Biol. 218, 559–579.

Suárez-Rivero, J., Villanueva-Paz, M., de la Cruz-Ojeda, P., de la Mata, M., Cotán, D., Oropesa-Ávila, M., de Lavera, I., Álvarez-Córdoba, M., Luzón-Hidalgo, R. and Sánchez-Alcázar, J. (2017). Mitochondrial Dynamics in Mitochondrial Diseases. Diseases 5, 1.

Vallat, R. (2018). Pingouin: statistics in Python. J. Open Source Softw. 3, 1026.

Warren, G. and Wickner, W. (1996). Organelle inheritance. Cell 84, 395–400.

Weisman, L. S. (2006). Organelles on the move: insights from yeast vacuole inheritance. Nat. Rev. Mol. Cell Biol. 7, 243–252.

Wu, H., Carvalho, P. and Voeltz, G. K. (2018). Here, there, and everywhere: The importance of ER membrane contact sites. Science 361,.

Yin, H., Pruyne, D., Huffaker, T. C. and Bretscher, A. (2000). Myosin V orientates the mitotic spindle in yeast. Nature 406, 1013–1015.

